# ERGA-BGE genome of *Androsace saussurei* Dentant, Lavergne, F. C. Boucher & S. Ibanez, a recently described plant from the rooftops of Europe

**DOI:** 10.1101/2025.01.21.634019

**Authors:** Sébastien Lavergne, Alexy Rosa, Chuan Peng, Rita Monteiro, Astrid Böhne, Thomas Marcussen, Torsten Struck, Rebekah A. Oomen, Genoscope Sequencing Team, Alice Moussy, Corinne Cruaud, Karine Labadie, Emilie Téodori, Lola Demirdjian, Patrick Wincker, Pedro H. Oliveira, Jean-Marc Aury, Tom Brown

## Abstract

The reference genome of *Androsace saussurei* (Ericales: Primulaceae: Primuloideae) provides a valuable resource for understanding the mechanisms of plant adaptations to the most extreme environments where flowering plants can thrive and the speciation mechanisms of plants dwelling on high alpine sky islands. The entirety of the genome sequence was assembled into 20 contiguous chromosomal pseudomolecules. This chromosome-level assembly encompasses 413 Mb, composed of 253 contigs and 101 scaffolds, with contig and scaffold N50 values of 4.95 Mb and 21.2 Mb, respectively.

## Introduction

*Androsace saussurei* is part of the plant family Primulaceae (Ericales) and is endemic to the highest summits of the Alps (Mont Blanc, Valais, Gran Paradiso, Vanoise) where it grows at the upper altitudinal limits of vascular plant life (from 2600 to 4300m asl in the Alps). The proposed species is one of three plant species recently described from the rooftops of Europe (Boucher et al. 2021). This species was named after the pioneer, alpine scientist, Horace Bénédict de Saussure (1740-1799), who first reported this plant in 1788 from the Mont Blanc range, but which remained unnamed until recent scientific explorations.

These plants form very rare and particular ‘cushion plant’ life forms that are extremely long-lived (several hundreds of years) and play a key role in plant and insect diversity at these elevations, acting as a shelter from other species. This role as a keystone engineer species is essential to the maintenance of multi-trophic diversity in high alpine conditions that are otherwise adverse to many species. The genus Androsace, encompassing ca 173 species (POWO, 2024), is typical of high alpine and arctic environments of the Northern Hemisphere, and hence the genome of the study species constitutes a valuable resource for studying the mechanisms and scenarios of plant speciation and adaptation to the most extreme environments where flowering plants can thrive.

The generation of this reference resource was coordinated by the European Reference Genome Atlas (ERGA) initiative’s Biodiversity Genomics Europe (BGE) project, supporting ERGA’s aims of promoting transnational cooperation to promote advances in the application of genomics technologies to protect and restore biodiversity (Mazzoni, Ciofi, and Waterhouse 2023).

## Materials & Methods

ERGA’s sequencing strategy includes Oxford Nanopore Technology (ONT) and/or Pacific Biosciences (PacBio) for long-read sequencing, along with Hi-C sequencing for chromosomal architecture, Illumina Paired-End (PE) for polishing (i.e. recommended for ONT-only assemblies), and RNA sequencing for transcriptomic profiling, to facilitate genome assembly and annotation.

### Sample and Sampling Information

Sébastien Lavergne, with help from C. Peng and A. Rosa, sampled one specimen of hermaphrodite *A saussurei*. The species was identified by Sébastien Lavergne based on the recent description of the species (Boucher et al., 2021), from Arète des Grandes Autannes, Le Tour, Chamonix, France on 7 July 2023. Sampling was performed under permit DDT-2023-1423 issued by the Préfecture de Haute Savoie (France). Sampling was performed via collection on the field, taken down through a one-hour hike, after which leaves and buds were picked from the plant and flash frozen using liquid nitrogen. Until DNA extraction, samples were preserved at -80C.

### Vouchering information

Physical reference material for the sequenced specimen has been deposited in Grenoble Museum, France, herbarium code Gr (https://www.grenoble.fr/74-museum-de-grenoble.htm) under the accession number GR005037.

Frozen reference tissue material of leaf from the same individual is available at the Biobank INRAE-CNRGV, Plant Genomic Center, Toulouse, France (https://cnrgv.toulouse.inrae.fr/fr) under the voucher ID 2024-PS3939.

### Data Availability

*Androsace saussurei* and the related genomic study were assigned to Tree of Life ID (ToLID) ddAndSaus6 and all sample, sequence, and assembly information are available under the umbrella BioProject PRJEB77232. The sample information is available at the following BioSample accessions: SAMEA114552830, and SAMEA115799864. The genome assembly is accessible from ENA under accession number GCA_964267185.1. Sequencing data produced as part of this project are available from ENA at the following accessions: ERX12733449, ERX12733450, and ERX12737191. Documentation related to the genome assembly and curation can be found in the ERGA Assembly Report (EAR) document available at https://github.com/ERGA-consortium/EARs/tree/main/Assembly_Reports/Androsace_saussurei/ddAndSaus6/. Further details and data about the project are hosted on the ERGA portal at https://portal.erga-biodiversity.eu/data_portal/Androsace%20saussurei.

### Genetic Information

The estimated genome size, based on ancestral taxa, is 1.67 Gb with a monoploid number of 10 chromosomes, a haploid number of 20 and a ploidy of four. All information for this species was retrieved from Genomes on a Tree (Challis et al., 2023).

We estimated the genome size and ploidy using PacBio HiFi data, analyzed with GenomeScope v2.0 and Smudgeplot v0.2.5, respectively. We report an estimated monoploid genome size of 237 Mb and a ploidy level of 4, tetraploid (Figure 1)

**Figure 1.**
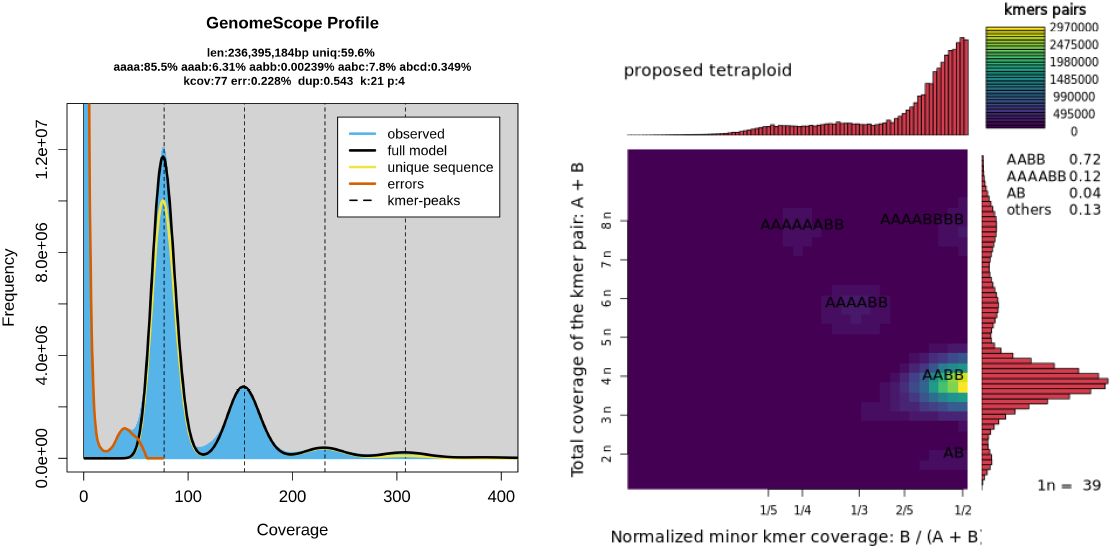
Kmer-based estimates of genome size, ploidy and heterozygosity. (Left) GenomeScope plot displaying the coverage of 31-mers (x-axis) against frequency x coverage of each 21-mer (y-axis). The black curve shows an inferred kmer distribution based on a ploidy of 4 and the inferred genome size (len), percentage of genome within homozygous (aaaa) or polymorphic regions (aaab, aabb, aabc, abcd), error rate (err) and unique sequence in the genome (uniq). The yellow curve shows the model based on unique 21-mers. The orange curve shows the curve based on the error 21-mers (Right) Smudgeplot displaying the normalised coverage (x-axis) against total coverage (y-axis) of each 21-mer in the sequencing dataset. Highlighted are elevated pairwise kmer coverage peaks, indicative of alternate ploidies. The probability of the best tested ploidy models are shown on the right, with the tetraploid model AABB showing the highest probability of 72%.

### DNA/RNA processing

DNA was extracted from 1 g of leaves using a conventional CTAB extraction followed by purification using Qiagen Genomic tips (QIAGEN, MD, USA). A detailed protocol is available on protocols.io (https://dx.doi.org/10.17504/protocols.io.bp2l694yzlqe/v1). DNA fragment size selection was performed using Short Read Eliminator (PacBio, CA, USA). Quantification was performed using a Qubit dsDNA HS Assay kit (Thermo Fisher Scientific) and integrity was assessed in a FemtoPulse system (Agilent). DNA was stored at 4 °C until usage.

RNA was extracted from leaves (50 mg) using the RNeasy Powerplant kit Plus Universal kit (Qiagen) following manufacturer instructions. Residual genomic DNA was removed with 6U of TURBO DNase (2 U/μL) (Thermo Fisher Scientific). Quantification was performed using a Qubit RNA HS Assay kit and integrity was assessed in a Bioanalyzer system (Agilent). RNA was stored at -80 °C.

### Library Preparation and Sequencing

Long-read DNA libraries were prepared with the SMRTbell prep kit 3.0 following manufacturers’ instructions and sequenced on a Revio system (PacBio). Omni-C libraries were prepared using the Dovetail Omni-C Kit (Dovetail Genomics, Scotts Valley, CA, USA) after plant nuclei isolation. Briefly, flash-frozen young leaves (1 g) were cryoground in liquid nitrogen and pure nuclei were first isolated following the “High Molecular Weight DNA Extraction from Recalcitrant Plant Species” protocol described by Workman et al. (https://doi.org/10.1038/protex.2018.059). Once the nuclei had been isolated, the pellets were treated as mammalian cells and Omni-C libraries were prepared according to the Mammalian Cell Protocol for Sample Preparation v1.4. The final libraries were sequenced on an Illumina NovaSeq 6000 instrument (Illumina, San Diego, CA, USA) with 2×150 bp read length. In total 76.5 Gb PacBio HiFi and 35.4 Gb HiC data were sequenced to generate the assembly.

### Genome Assembly Methods

The genome of *Androsace saussurei* was assembled using NextDenovo v2.5.1, followed by scaffolding using YaHS v1.2. The assembled scaffolds were then curated through manual inspection using PretextView v0.2.5 to remove any false joins and to incorporate sequences not automatically scaffolded into their respective locations within the chromosomal pseudomolecules. The mitochondrial and chloroplast genomes were assembled as single circular contigs using Oatk v1.0 and included in the released assembly (GCA_964267185.2).

As the genome is tetraploid, allelic duplications were not removed after the initial assembly. Instead, the two haplotypes were separated during manual curation, resulting in two chromosome-scale assemblies, each comprising 20 chromosomes (GCA_964267185.2 and GCA_964267195.1). Summary analysis of the released assembly was performed using the ERGA-BGE Genome Report ASM Galaxy workflow (https://doi.org/10.48546/workflowhub.workflow.1104.1).

## Results

### Genome Assembly

The primary genome assembly has a total length of 413,403,326 bp in 101 scaffolds including the mitochondrial and chloroplast genomes (Figures 2 & 3), with a GC content of 33.05%. The assembly has a contig N50 of 4,952,245 bp and L50 of 29 and a scaffold N50 of 21,172,409 bp and L50 of 10 with a total of 152 gaps. The single-copy gene content analysis using the Eudicots database with BUSCO (Manni et al. 2021) resulted in 95.0% completeness (58.0% single and 37.0% duplicated). 63.3% of reads k-mers were present in the assembly and the assembly has a base accuracy Quality Value (QV) of 53.6 as calculated by Merqury (Rhie et al. 2020).

**Figure 2.**
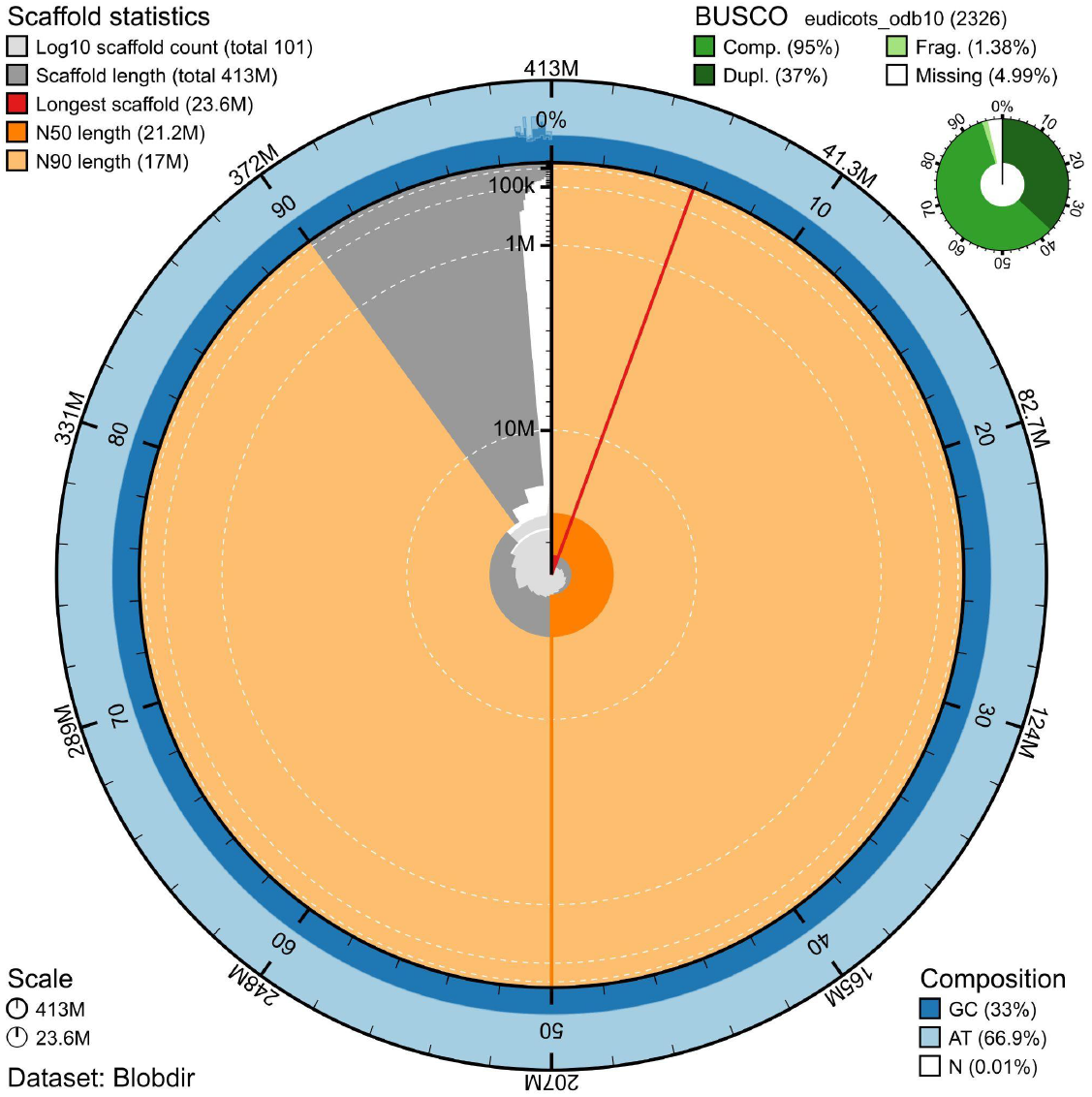
Snail plot summary of assembly statistics. The main plot is divided into 1,000 size-ordered bins around the circumference, with each bin representing 0.1% of the 413,403,326 bp assembly including the mitochondrial genome. The distribution of sequence lengths is shown in dark grey, with the plot radius scaled to the longest sequence present in the assembly (23,550,734 bp, shown in red). Orange and pale-orange arcs show the scaffold N50 and N90 sequence lengths (21,172,409 and 17,031,950 bp), respectively. The pale grey spiral shows the cumulative sequence count on a log-scale, with white scale lines showing successive orders of magnitude. The blue and pale-blue area around the outside of the plot shows the distribution of GC, AT, and N percentages in the same bins as the inner plot. A summary of complete, fragmented, duplicated, and missing BUSCO genes found in the assembled genome from the Eudicots database (odb10) is shown in the top right.

**Figure 3.**
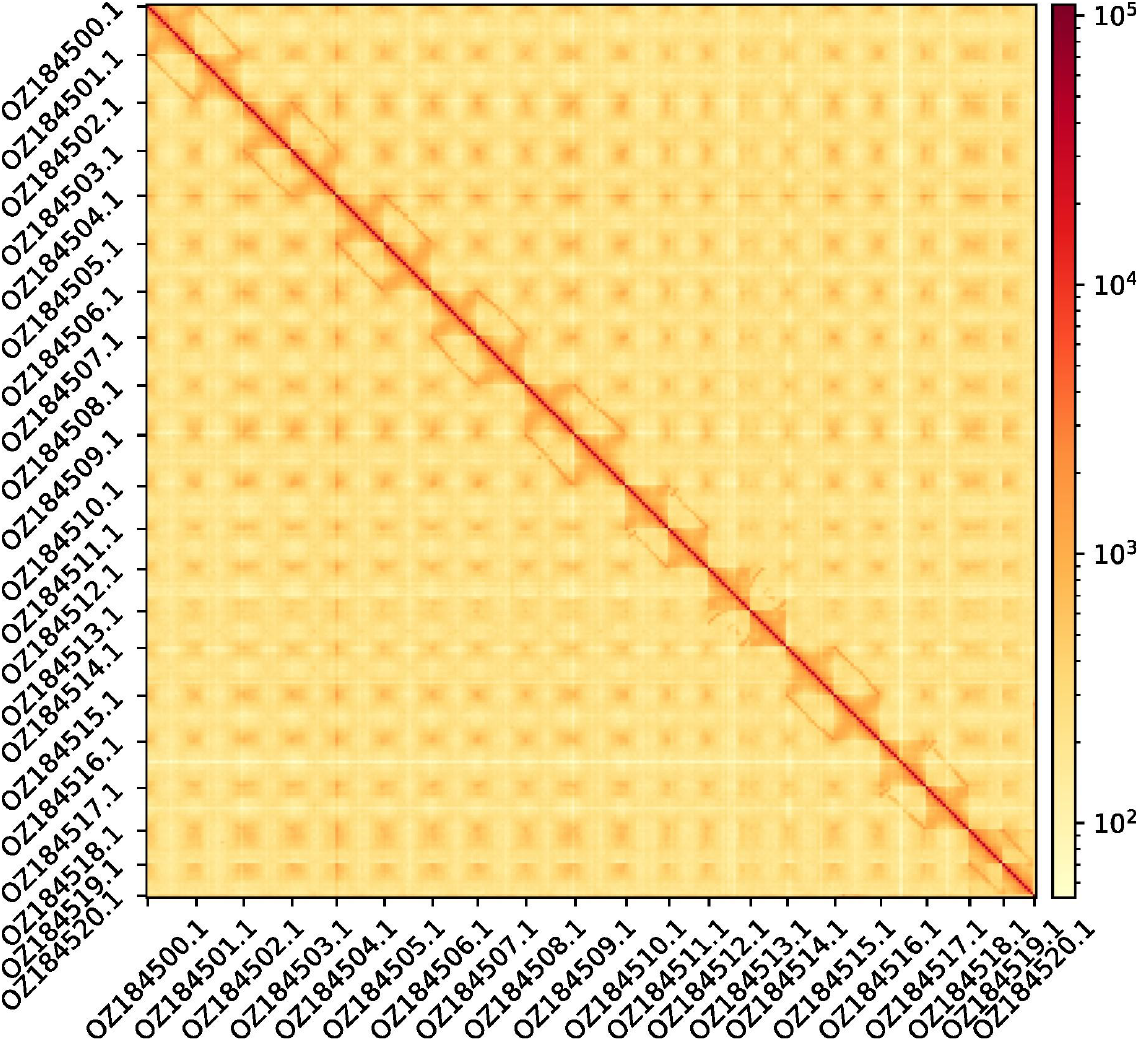
Hi-C contact map showing spatial interactions between regions of the genome. The diagonal corresponds to intra-chromosomal contacts, depicting chromosome boundaries. The frequency of contacts is shown on a logarithmic heatmap scale. Hi-C matrix bins were merged into a 25 kb bin size for plotting.

## Acknowledgements

We acknowledge the co-descriptors of the species, Cédric Dentant, Florian Boucher and Sébastien Ibanez. We would like to acknowledge the assembly reviewer, Sarah Pelan, from the Wellcome Sanger Institute. The authors acknowledge the support of the Freiburg Galaxy Team: Saim Momin and Björn Grüning, Bioinformatics, University of Freiburg (Germany), funded by the German Federal Ministry of Education and Research BMBF grant 031 A538A de.NBI-RBC and the Ministry of Science, Research and the Arts Baden-Württemberg (MWK) within the framework of LIBIS/de.NBI Freiburg.

## Author Contributions

SL, CP and AR collected the species, SL identified the species, SL sampled and preserved biological material and provided metadata, TM, RM, RO, THS and AsB provided support in sampling, shipping of biological material, metadata collection, and management, the GST extracted DNA, prepared libraries, and performed sequencing under the supervision of AM, CC, KL, PHO and PW; LD, JMA and ET performed genome assembly and curation, TB generated the analysis and report. All authors contributed to the writing, review, and editing of this genome note and read and approved the final version. This work is part of the species assigned to Genoscope, which was instrumental in the wet lab, sequencing, and assembly processes, and represents a key contribution to BGE’s outputs

## Author Information

Members of the Genoscope Sequencing Technical Team are listed here: https://doi.org/10.5281/zenodo.14611490

## Literature Cited

Boucher, Florian C., Cédric Dentant, Sébastien Ibanez, Thibaut Capblancq, Martí Boleda, Louise Boulangeat, Jan Smyčka, Cristina Roquet, and Sébastien Lavergne. 2021. “Discovery of Cryptic Plant Diversity on the Rooftops of the Alps.” Scientific Reports 11 (1): 11128.

Manni, Mosè, Matthew R. Berkeley, Mathieu Seppey, Felipe A. Simão, and Evgeny M. Zdobnov. 2021. “BUSCO Update: Novel and Streamlined Workflows along with Broader and Deeper Phylogenetic Coverage for Scoring of Eukaryotic, Prokaryotic, and Viral Genomes.” Molecular Biology and Evolution 38 (10): 4647–54.

Mazzoni, Camila J., Claudio Ciofi, and Robert M. Waterhouse. 2023. “Biodiversity: An Atlas of European Reference Genomes.” Nature 619 (7969): 252.

Rhie, Arang, Brian P. Walenz, Sergey Koren, and Adam M. Phillippy. 2020. “Merqury: Reference-Free Quality, Completeness, and Phasing Assessment for Genome Assemblies.” Genome Biology 21 (1): 245.

POWO (2024). Plants of the World Online. Facilitated by the Royal Botanic Gardens, Kew. Published on the Internet; https://powo.science.kew.org/ Retrieved 05 December 2024.

